# Silicon supplementation and jasmonate activation synergistically increase phenolic defences against a legume herbivore

**DOI:** 10.1101/2023.03.01.530676

**Authors:** J.M.W. Ryalls, A.N. Gherlenda, R.C. Rowe, B.D. Moore, S.N. Johnson

## Abstract

1. The accumulation of silicon (Si) is widely reported to have anti-herbivore defensive properties in grasses. There is emerging, but fragmentary, evidence that Si could play a similar role in legumes.
2. Here, we sought to understand the effects of Si supplementation on anti-herbivore defensive properties in lucerne (*Medicago sativa*), especially in relation to other potential defences (i.e. phenolics) and the phytohormone that regulates anti-herbivore defences, jasmonic acid or jasmonate (JA), which is also linked to Si accumulation.
3. We determined how growth, root nodulation and chemistry (carbon, nitrogen and phenolic concentrations) of four genotypes of lucerne responded to Si supplementation, with and without the application of JA, and we used feeding assays to determine the subsequent effects on the feeding success of adult *Sitona discoideus* weevils.
4. Si supplementation increased plant mass and root nodulation of *M. sativa* by 61% and 227%, respectively, and reduced relative consumption (RC) and frass production by *S. discoideus* by 38% and 30%, respectively. Si supplementation had no effect on foliar nitrogen concentrations, most likely due to the dilution effects of increased plant growth and foliar carbon. Phenolic concentrations were negatively correlated with leaf RC; RC also decreased by 34% when JA was applied to plants. When Si was combined with JA application, phenolics were significantly enhanced, demonstrating the potential to stimulate multiple anti-herbivore properties in *M. sativa*.
5. **Synthesis**. The novel findings suggest that Si accumulation may play a more important role in legume resistance to herbivorous animals than previously thought. The ubiquity of soil Si and its emerging functional role in plant biology, including plant–animal interactions, suggest that these patterns could be common amongst legumes.

## Introduction

Silicon (Si) accumulation by plants is now recognised as a major mechanism for alleviating the adverse effects of environmental stresses, including drought, metal toxicity, disease and herbivory (Cooke & Leishman 2016; Debona, Rodrigues & Datnoff 2017). Si is taken up by roots as monosilicic acid, Si(OH)_4_, and subsequently translocated and deposited in various plant tissues, within and between cells, in the form of biogenic silica, SiO_2_ (Raven 1983). Si defences against herbivores include abrasive effects on their mouthparts and diminished nutrient acquisition via reduced palatability and digestibility of foliage (Massey & Hartley 2006; Hartley & DeGabriel 2016).

The vast majority of research into the beneficial effects of Si supplementation of plants is concerned with the grasses (Poaceae) due to many of them being hyper-accumulators of Si (Massey, Ennos & Hartley 2006; Johnson *et al*. 2021). A recent review, however, revealed that Si supplementation of legumes (Fabaceae) alleviated similar environmental stresses as grasses (Putra *et al*. 2020). Legumes are the most valuable source of protein for crops and animal feed, and also provide important habitats for beneficial organisms (e.g. pollinators) (Stagnari *et al*. 2017). A key knowledge gap that Putra *et al*. (2020) identified was how little is known about whether Si alleviated the impacts of herbivory in legumes. To our knowledge, only one study has investigated the impacts of Si supplementation of a legume (soybean; *Glycine max*) on a chewing insect herbivore, the feeding guild most affected by Si defences (Johnson *et al*. 2021), which reported reduced herbivore growth rates (Johnson, Rowe & Hall 2020). Cell– and sap–feeding arthropods have similarly been negatively impacted by Si supplementation of soybean (Ferreira & Moraes 2011) and the common bean (*Phaseolus vulgaris*) (Peixoto *et al*. 2011).

In addition to providing physical defence against herbivores, Si supplementation is known to alter the activity of the jasmonate (jasmonic acid (JA)) pathway (Hall *et al*. 2019). JA is the master regulator of secondary metabolite defences (e.g. phenolics) against chewing arthropods (Puentes *et al*. 2021). Stimulation of the JA pathway also causes rapid induction of Si defences (Ye *et al*. 2013; Johnson *et al*. 2021; Waterman *et al*. 2021). Si accumulation in grasses tends to trade off with phenolic concentrations because Si is used as a metabolically cheaper defence against herbivores (Cooke & Leishman 2012). Legumes accumulate relatively small concentrations of Si compared with the grasses (Putra *et al*. 2020), so it is possible that that this trade-off is less apparent if present at all. Moreover, activation of the JA pathway decoupled the negative relationship between Si and phenolic defences in the grass *Brachypodium distachyon* (Waterman *et al*. 2021) so the relationship between Si accumulation and phenolics is somewhat plastic.

Legumes possess root nodules that house nitrogen (N)-fixing rhizobial bacteria, which increase nitrogen availability for legumes and subsequent or adjacent non-N-fixing plants (Chmelíková *et al*. 2015; Ryalls *et al*. 2016). Si supplementation frequently increases nodule abundance, nitrogenase activity (indicative of increased N-fixation) and concentrations of nitrogenous compounds (protein and amino acids) in *Medicago* spp. and other legumes (Nelwamondo & Dakora 1999; Izaguirre-Mayoral *et al*. 2017; Putra *et al*. 2022), especially when plants are under stress (e.g. Al Murad & Muneer 2022). This could potentially undermine the effectiveness of Si in increasing legume resistance to herbivory because N is usually the limiting nutrient in insect herbivore diets (Mattson 1980); increased N availability via Si supplementation could therefore increase susceptibility to herbivory, assuming that this increase in N is not outweighed by the Si-induced reduction in digestibility of that N.

It could be envisaged that Si supplementation of legumes could affect herbivorous insects in a multitude of ways, which we conceptually depict in Fig. 1. In summary, Si supply may promote leaf silicification and Si-based defences in the leaves (mechanism 1; Fig. 1) and could either increase or decrease concentrations of secondary metabolite defences such as phenolics (mechanism 2; Fig. 1). Concurrently, stimulation of the JA pathway may promote Si accumulation and phenolic defences (mechanism 3; Fig. 1), both of which may negatively affect herbivore feeding and performance. An added consideration for legumes, compared with non-leguminous plants, is that Si supplementation may also make plants more nutritious to herbivores by increasing N concentrations in leaves (mechanism 4; Fig. 1).

**Fig. 1.**
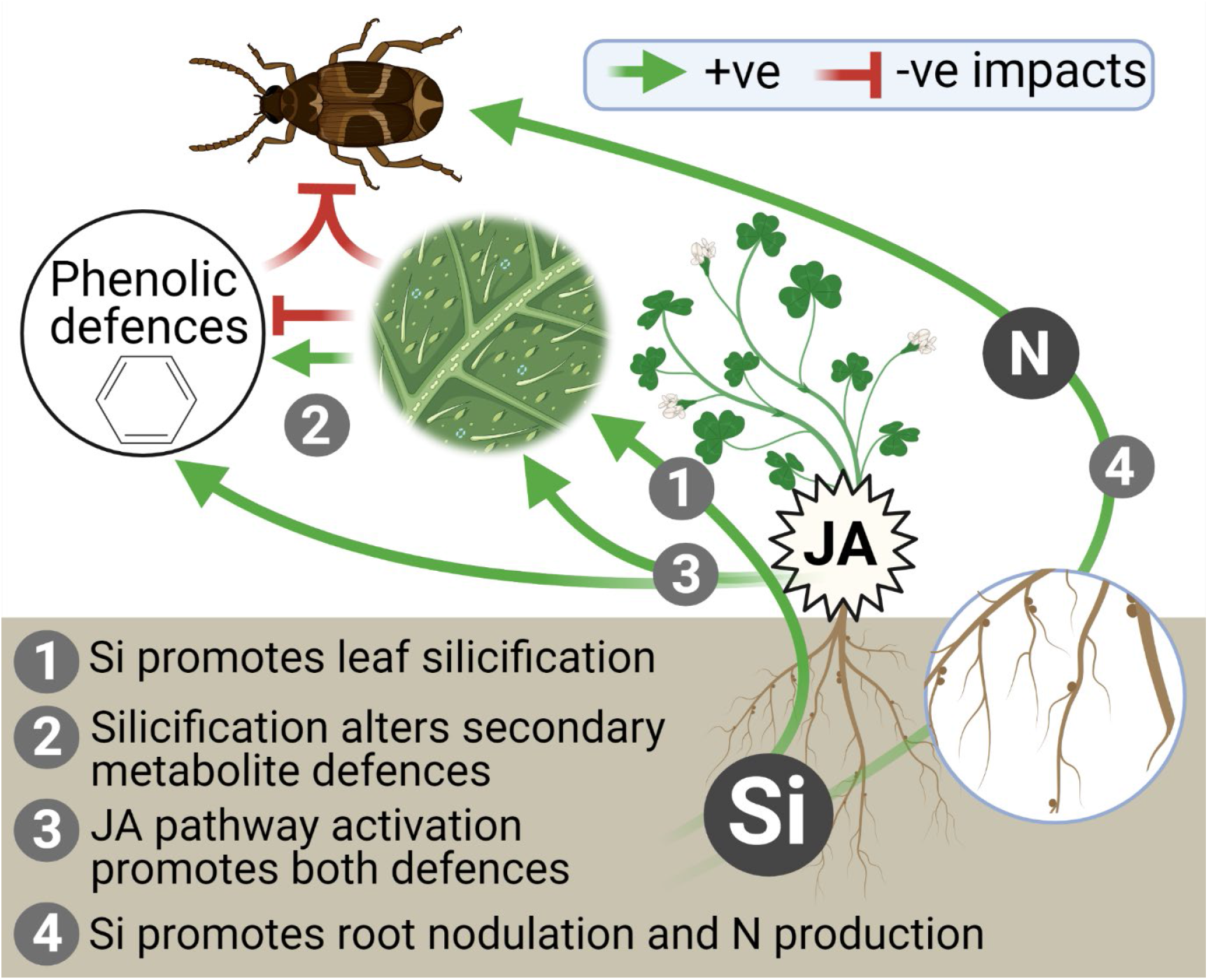
Conceptual model, based on the available literature, for how Si supplementation and JA application may affect Si and phenolic defences against insect herbivores in legumes.

The aim of this study was to better understand the role of Si in anti-herbivore defences in a legume, particularly in relation to other defences (i.e. phenolics) and the JA pathway. We investigated the effects of both Si supplementation and JA application on four genotypes of lucerne (also known as alfalfa), *Medicago sativa*. We quantified changes in foliar concentrations of Si, carbon (C), N and phenolics and determined how this affected feeding behaviour and growth rates of a chewing folivore (adult *Sitona discoideus* weevils). As summarised in Fig. 1, we hypothesised that:

1. Si supplementation increases foliar concentrations of Si (silicification),
2. Foliar silicification alters concentrations of phenolics; both defences reduce feeding activity and performance of *S. discoideus*,
3. Stimulation of the JA pathway increases both Si and phenolic defences,
4. Si supplementation increases foliar concentrations of N which may moderate defences against *S. discoideus*.

## Materials and methods

### Plant and insect material

Thirty two plants of each of four lucerne genotypes (Hunter River, Sequel, Trifecta and Genesis; as described in Ryalls *et al*. (2017)) were grown from seed (128 plants in total) in 70 mm pots filled with sieved (2mm) local loamy-sand soil collected from the Hawkesbury Forest Experiment in Richmond, NSW (see Ryalls, Moore & Johnson 2018 for chemical composition). Plants were maintained in growth chambers at 26/18 °C day/night on a 15L:9D cycle for 12 weeks (February-May 2017). Humidity was controlled at 60%. Half of the plants were supplemented with Si (500 mg l^-1^ solution of dissolved sodium metasilicate) and half received deionized water alone (c. 23 mL three times a week in both cases), as in Ryalls *et al*. (2018). Three days before the feeding assay, half of the Si-supplemented plants were treated with methyl jasmonate (MeJA; 95%, Sigma Aldrich) by applying 1 mL of a solution of 100 μg mL^-1^ MeJA in Tween 20 ± 0.1% to the base of the stem at the surface of the soil with a pipette, as in Johnson *et al*. (2018a). The other half were supplemented with 1 mL of Tween 20 only. This was repeated 24 and 48 h later. 160 sexually mature *S. discoideus* adults were collected from a local lucerne field in Richmond, NSW and maintained on propagated lucerne plants (40 weevils on each of the four genotypes) under the same conditions as the plants for one week prior to the feeding assay.

### Experimental procedure – Feeding assay

Four leaves were removed from each plant. The two youngest leaves remained on the plant and were not used. Two were weighed (using a 0.001 mg micro-balance) and placed in an empty Petri dish and the other two were weighed and placed in a Petri dish containing one adult weevil, which was starved for 24 h and weighed prior to the feeding assay. After 24 h, remaining leaves were dried and reweighed. The dry mass eaten was calculated by subtracting the dry mass after feeding from the dry mass before feeding [i.e. fresh weight offered × (dry mass control at end/wet mass control at start)]. This was used to calculate the relative consumption (RC) for each adult weevil (equation 1).

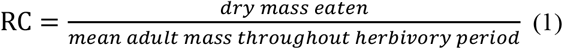

Weevils were weighed after the feeding assay and starved for a further 12 h to ensure all frass was expelled, before being reweighed. Relative growth rate (RGR) was also defined for each adult weevil (equation 2).

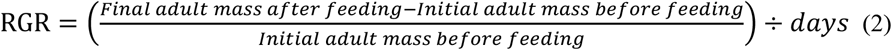

Once the feeding assay had been initiated, remaining aboveground and belowground plant parts were removed from the soil. Remaining soil was washed from plants over 200 μm and 20 μm stacked sieves and the numbers of nodules on the roots were counted. Roots were separated from shoots and all plant material was snap-frozen in liquid N before being freeze-dried and weighed.

### Chemical analyses

Dried leaf material was separated from stems and ball-milled to a fine powder prior to chemical analyses. Total Si concentrations (% of dry mass) were determined from all 128 plants with an X-ray fluorescence spectrometer (Epsilon-3x, PANalytical, EA Almelo, The Netherlands) using small mass holders, following the protocol described by Hiltpold *et al*. (2016). Si data were calibrated using citrus plant material of known Si concentration. Carbon (C) and nitrogen (N) concentrations were determined for half of the leaf samples (64 plants; N = 4 per genotype per Si treatment per JA treatment) with a Carlo Erba CE1110 elemental analyser. Total phenolics were quantified from the other half of the plants (N = 64) using the Prussian blue assay and gallic acid as a standard (Graham 1992), following the protocol described by Waterman *et al*. (2021). Absorbance was averaged from three replicate extracts per sample and total phenolic concentration (TPC; mg GAE/g dry weight) was calculated using the formula TPC = cV/m, where c = concentration of gallic acid obtained from a calibration curve of known standards in mg/mL, V = volume of extract (0.5 mL), and m = mass of extract (0.05g).

### Statistical analyses

The R statistical interface v4.2.0 was used for all statistical analyses. The effects of genotype, Si supplementation and JA application on plant growth (root mass, shoot mass and nodulation), foliar chemistry (%Si, %C, %N and total phenolics) and insect feeding metrics (RC, frass production and RGR) were analysed using general linear models. The fixed effects for all models included genotype (Aurora, Genesis, Sequel and Trifecta), Si treatment (Si- and Si+) and JA treatment (JA-and JA+) as well as the two-way and three-way interactions between these terms. *Post-hoc* tests using the package *emmeans* were used for pairwise comparisons of means. Where appropriate, dependent variables were transformed before analysis (Table 1). Pearson’s correlations were used to determine whether C-based defence (phenolics) was associated with RC, frass production or RGR.

**Table 1.** Plant growth, foliar chemistry and weevil responses to plant genotype, Si treatment (Silicon) and jasmonic acid (JA) treatment from linear models.

| Response variable | Fig. | df | Genotype |  | Silicon |  | JA |  | Genotype<br>× Silicon |  | Genotype<br>× JA |  | Silicon<br>× JA |  | Genotype<br>× Silicon<br>× JA |  |
| --- | --- | --- | --- | --- | --- | --- | --- | --- | --- | --- | --- | --- | --- | --- | --- | --- |
|  |  |  | *F*/*χ*<sup>2</sup><sub>3</sub> | *P* | *F*/*χ*<sup>2</sup><sub>1</sub> | *P* | *F*/*χ*<sup>2</sup><sub>1</sub> | *P* | *F*/*χ*<sup>2</sup><sub>3</sub> | *P* | *F*/*χ*<sup>2</sup><sub>3</sub> | *P* | *F*/*χ*<sup>2</sup><sub>1</sub> | *P* | *F*/*χ*<sup>2</sup><sub>3</sub> | *P* |
| Plant growth |  |  |  |  |  |  |  |  |  |  |  |  |  |  |  |  |
| Shoot mass (mg)<sup>a</sup> | 2A | 112 | 3.47 | **0.019** | 17.68 | < **0.001** | 0.36 | 0.55 | 0.98 | 0.404 | 0.58 | 0.628 | 0.11 | 0.741 | 0.16 | 0.921 |
| Root mass (mg)<sup>a</sup> | 2A | 112 | 10.76 | < **0.001** | 29.82 | < **0.001** | 2.29 | 0.133 | 1.26 | 0.293 | 0.33 | 0.802 | 0.06 | 0.803 | 0.75 | 0.526 |
| Nodulation (#) | 2B | 112 | 8.12 | **0.044** | 22.89 | < **0.001** | 0.01 | 0.903 | 4.13 | 0.248 | 0.33 | 0.954 | 1.72 | 0.190 | 3.51 | 0.320 |
| Foliar chemistry |  |  |  |  |  |  |  |  |  |  |  |  |  |  |  |  |
| Silicon (%)<sup>a</sup> | 3A | 112 | 0.40 | 0.758 | 14.75 | < **0.001** | 0.62 | 0.433 | 0.16 | 0.925 | 0.55 | 0.646 | 0.03 | 0.870 | 0.35 | 0.791 |
| Carbon (%) | 3B | 48 | 0.68 | 0.570 | 4.44 | **0.040** | 6.73 | **0.013** | 1.25 | 0.303 | 0.72 | 0.543 | 0.14 | 0.708 | 1.42 | 0.248 |
| Nitrogen (%) | 3C | 48 | 1.71 | 0.178 | 0.03 | 0.860 | 0.95 | 0.333 | 0.49 | 0.690 | 0.29 | 0.835 | 0.51 | 0.480 | 0.05 | 0.986 |
| TPC (mg/g DW) | 3D | 48 | 0.86 | 0.469 | 4.06 | **0.049** | 39.39 | < **0.001** | 0.43 | 0.734 | 0.04 | 0.990 | 4.96 | **0.031** | 0.05 | 0.987 |
| Weevil feeding metrics |  |  |  |  |  |  |  |  |  |  |  |  |  |  |  |  |
| RC | 4A | 111 | 2.36 | 0.076 | 15.96 | < **0.001** | 12.16 | < **0.001** | 1.11 | 0.347 | 0.14 | 0.934 | 0.34 | 0.562 | 0.10 | 0.959 |
| Frass (mg)<sup>a</sup> | 4B | 110 | 1.35 | 0.261 | 12.98 | < **0.001** | 1.85 | 0.176 | 0.14 | 0.937 | 0.61 | 0.613 | 0.05 | 0.822 | 0.40 | 0.754 |
| RGR (mg/mg/day) | S1 | 111 | 3.42 | **0.020** | 1.05 | 0.307 | 0.91 | 0.343 | 0.70 | 0.554 | 1.81 | 0.149 | 1.38 | 0.243 | 2.01 | 0.117 |

P-values highlighted in bold indicate significance ( $P < 0.05$ ). Where appropriate, response variables were transformed (<sup>a</sup>square root, <sup>b</sup>log) before analysis. TPC, RC and RGR refer to total phenolic concentration, relative consumption and relative growth rate, respectively. Nodulation (negative binomial model) is reported as a likelihood-ratio $\chi^2$ test statistic. All other parameters (general linear models) are reported as F-values.

## Results

### Plant growth

Si supplementation significantly increased shoot mass (Fig. 2a), root mass (Fig. 2a) and the number of root nodules (Fig. 2b). Full statistical results are shown in Table 1. Root mass, shoot mass and nodulation tended to be lowest in the genotype Aurora compared with the three other genotypes (Fig. 2). No significant effects of JA treatment, and no interactions between fixed effects were observed.

**Fig. 2.**
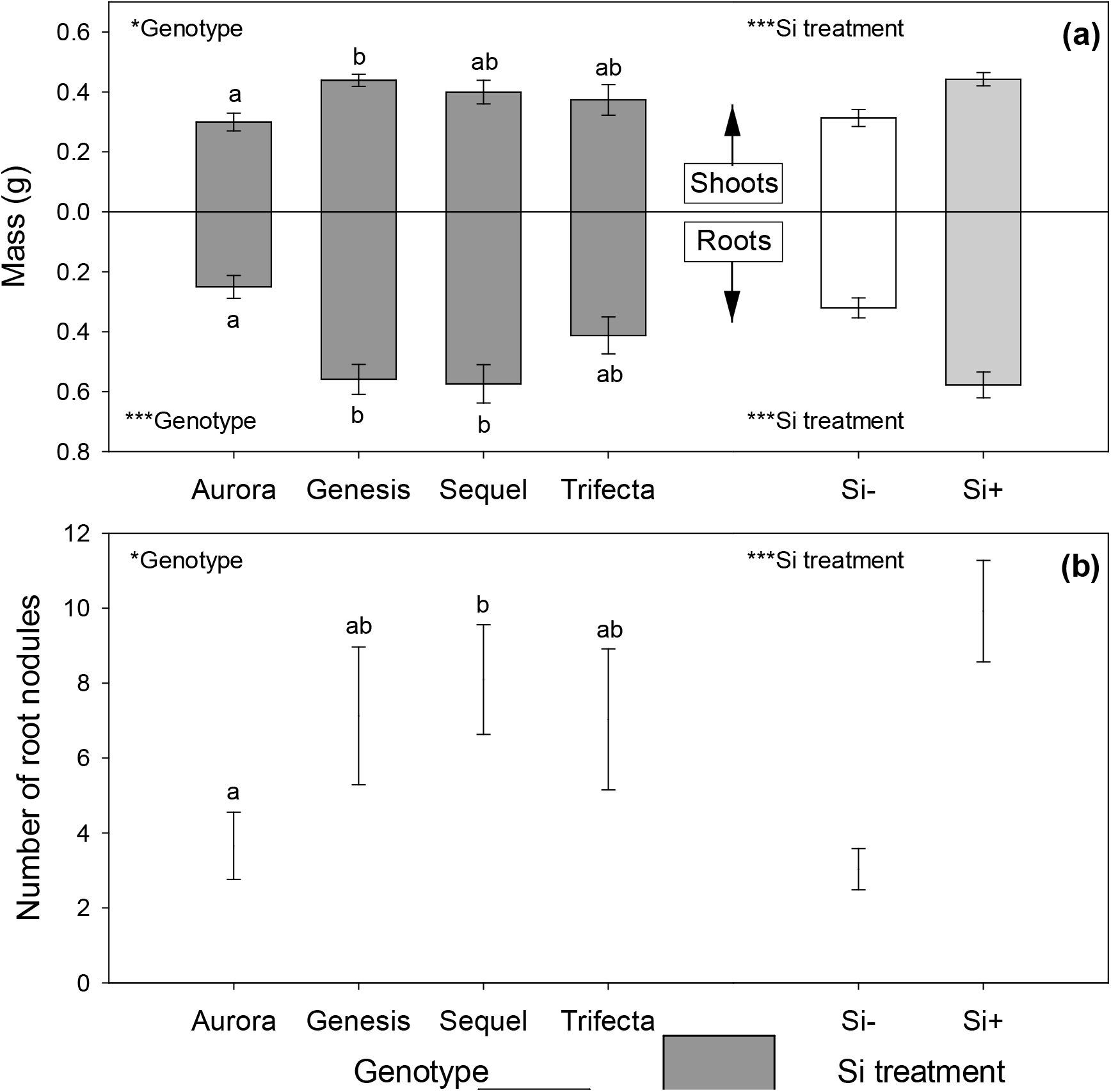
The effects of genotype and silicon (Si) supplementation on the shoot and root mass (a) and root nodulation (b) of *Medicago sativa* plants. Values are means ± SE. Bars with the same letters were not significantly different (*P* < 0.05). Statistical significance of fixed effects indicated by *(0.01 < *P* < 0.05) and ***(*P* < 0.001).

### Foliar chemistry

Si supplementation significantly increased foliar Si (Fig. 3a) and C concentrations (Fig. 3b). JA treatment also increased the concentration of foliar C (Fig. 3b). Foliar N concentrations were not affected by Si or JA treatments (Fig. 3c), individually or in combination. Si treatment in interaction with JA treatment significantly affected total phenolics, which increased when JA was applied, especially when plants were supplemented with Si (Fig. 3d). Genotype had no significant effect on any of the foliar chemicals measured (Table 1).

**Fig. 3.**
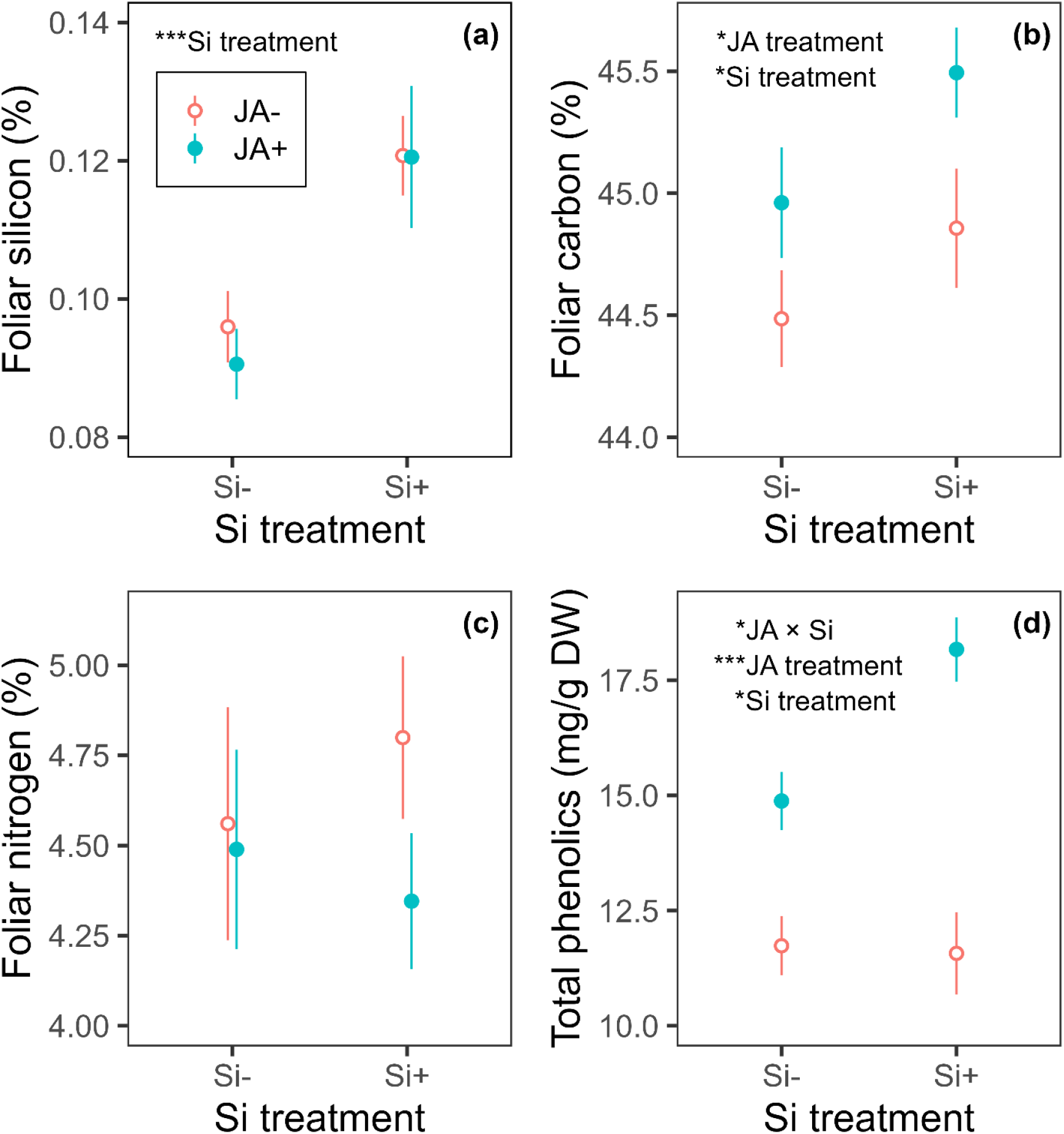
The effects of jasmonic acid (JA) application and silicon (Si) supplementation on foliar silicon (a), carbon (b), nitrogen (c) and total phenolic concentrations (d) of *Medicago sativa*. Values are means ± SE. Statistical significance of fixed effects indicated by *(0.01 < *P* < 0.05) and ***(*P* < 0.001).

### Weevil feeding metrics

Si supplementation significantly decreased both the RC of leaf material (Fig. 4a) and the amount of frass produced (Fig. 4b). While JA application significantly decreased RC (Fig. 4a), frass production was not significantly lower when JA was applied (Fig. 4b). No genotype effect, two-way or three-way interactive effects on RC or frass production were observed. RGR was not significantly affected by JA or Si treatments, individually or interactively. However, RGR was significantly higher in the genotype Genesis compared with Sequel (Figure S1). Total phenolic concentration was significantly correlated with RC (increasing phenolics decreased RC; r = - 0.259, *P* = 0.041, df = 61), but not frass production (r = -0.200, *P* = 0.116, df = 61) or RGR (r = - 0.001, *P* = 0.974, df = 61).

**Fig. 4.**
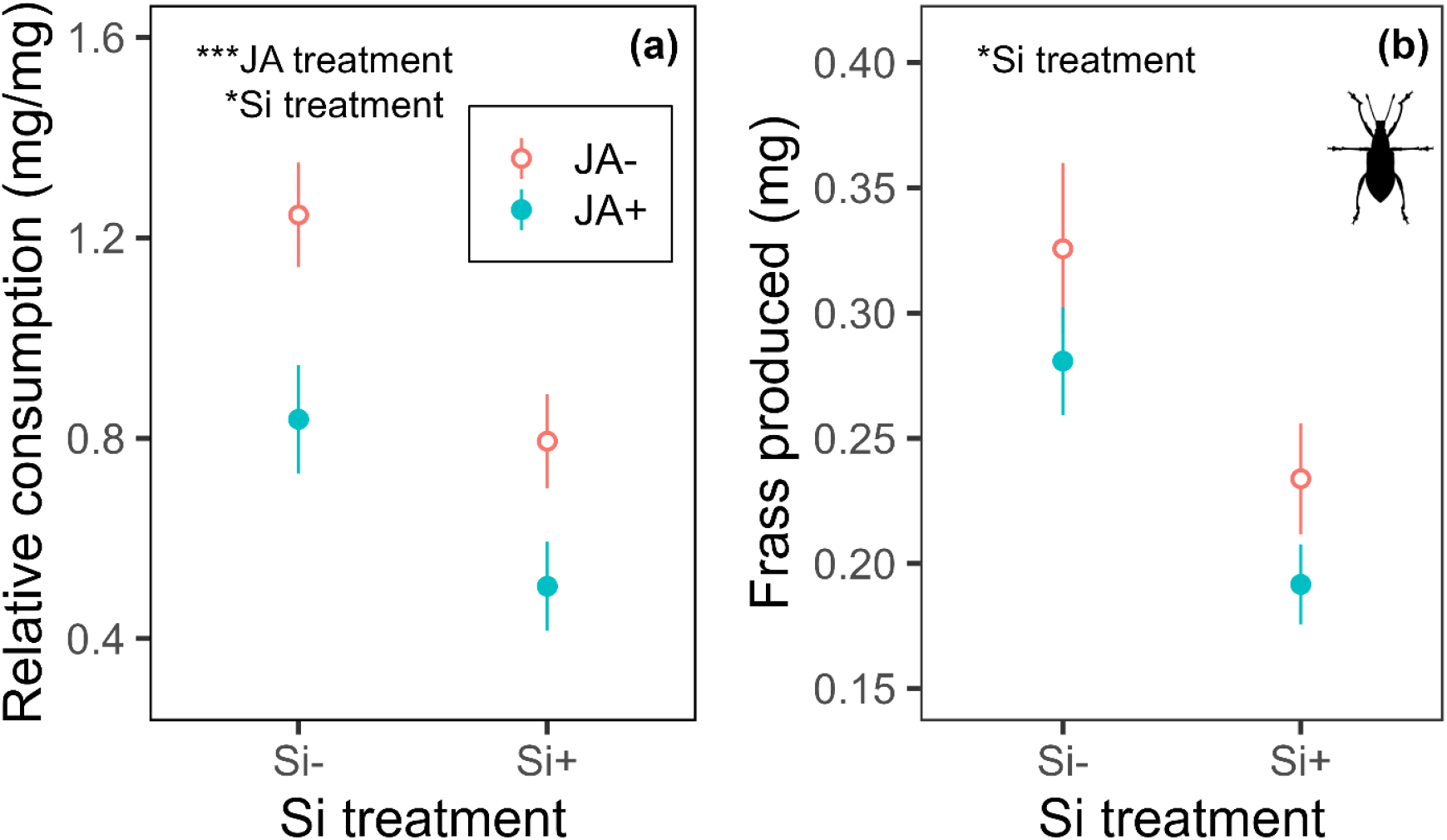
The effects of jasmonic acid (JA) application and silicon (Si) supplementation on relative consumption (a) and frass production (b) by *Sitona discoideus*. Values are means ± SE. Statistical significance of fixed effects indicated by *(0.01 < *P* < 0.05) and ***(*P* < 0.001). Weevil image by JCGiron using the R package *phylopic*.

## Discussion

### Si supplementation increased foliar Si in lucerne

Most studies on biotic responses to plant Si deposition have used species of Poaceae (mostly cereal crops) as high Si-accumulating model plants (Cooke & Leishman 2016; Johnson *et al*. 2021), in which Si is assumed to be more functionally important. Foliar Si concentrations in legumes have been shown to increase (Johnson, Rowe & Hall 2020), decrease (Johnson *et al*. 2018b) or remain unchanged (Johnson *et al*. 2017) in response to Si supplementation and the outcome may depend on the diversion of resources towards plant growth and nodule development (Putra *et al*. 2022).

### Si supplementation promoted accumulation of phenolics

While previous studies have identified trade-offs between foliar Si and C-based defences such as phenolics (Cooke & Leishman 2012; Johnson & Hartley 2018), we observed the opposite in *M. sativa* in this current study. Other studies have suggested that Si potentially optimises other plant defences as part of a wider herbivore defensive syndrome (Moles *et al*. 2013; Cibils-Stewart *et al*. 2022). Si deposits in tissues may exacerbate the physical aggravation caused by herbivore damage to increase JA activity and downstream production of secondary metabolite defences (Hall *et al*. 2019). This may therefore explain why Si supplementation promoted the production of phenolics when JA was applied, since stimulation with MeJA simulates herbivore attack. In the absence of this stimulation, foliar phenolic concentrations were largely unchanged by Si supplementation.

### Stimulation of the JA pathway stimulated phenolic production and reduced herbivore feeding

We observed a JA-induced reduction in relative consumption by *S. discoideus* weevils, which was associated with an increase in phenolics. Phenolics and relative consumption were negatively correlated, although this trend was not reflected in the short-term growth of *S. discoideus*. The application of JA has also promoted phenolic production in other plant species, such as *Catharanthus roseus* (Liu *et al*. 2016), *Brassica oleracea* (Guan *et al*. 2019), *Ocimum basilicum* (Malekpoor, Salimi & Pirbalouti 2016) and *Picea abies*, this latter example been associated with resistance to the pine weevil, *Hylobius abietis* (Puentes *et al*. 2021). While consuming less food tends to decrease herbivore frass production, decreased digestibility of food can counteract this (Massey & Hartley 2009), which may explain why JA supplementation and phenolic production did not significantly alter the frass production of *S. discoideus*.

### Si increased root nodulation but not foliar N concentrations

Root nodule abundance increased in response to Si supplementation, which has previously been demonstrated in *M. sativa* (Johnson *et al*. 2017; Johnson *et al*. 2018b) and *M. truncatula* (Putra *et al*. 2022). This may have been a compensatory response to higher plant growth (and its associated increase in foliar C concentrations) to prevent the plant from becoming N-limited. Alternatively, Si-induced root nodulation may have provided the N, allowing for increased growth. However, this Si-induced nodulation response was not reflected in foliar N concentrations. Studies using grass species tend to show negative effects of Si supplementation on foliar N concentrations, whereas this relationship is likely to be less apparent in legumes, which only accumulate low to moderate levels of Si (Putra *et al*. 2021). Si has been shown to promote foliar N and N-containing compounds in *Medicago* spp. (Johnson *et al*. 2018b; Putra *et al*. 2022). In this study, however, increased plant growth may have caused a dilution effect, whereby increased uptake of foliar N was accompanied by greater tissue mass. Moreover, root Si deposition may anatomically alter the structure of root nodules but does not necessarily promote rhizobial N-fixation (Putra *et al*. 2021) and foliar N concentrations may be dictated by rhizobial efficacy. For example, when low-efficiency rhizobial strains were present in *M. truncatula*, foliar N was found to increase, whereas plants with high-efficiency strains had no effect on foliar N concentrations (Putra *et al*. 2022).

### Conclusions

Given the importance of N-fixation in legumes to ecosystem function and the potential for Si to positively impact the symbiosis between legumes and rhizobia bacteria in root nodules, surprisingly few studies have examined the effects of Si on legumes. Si clearly has the potential to increase nodulation and plant growth of legumes, while reducing herbivore feeding. Moreover, when Si is combined with an exogenous application of JA, it has the potential to stimulate additional anti-herbivore defence. As such, the field of insect–plant ecology would benefit from further understanding into Si dynamics in legumes and their responses to JA application to enhance crop protection and ecosystem productivity.

## Acknowledgements

This research was funded by a Discovery Project grant from the Australian Research Council (ARC DP140100636) awarded to SNJ and BDM. JMWR is funded by a Leverhulme Trust Early Career Fellowship (ECF-2020-017). We would like to thank Dr Lisa Bromfield for their help with mist netting to collect weevils.

## Author contributions

JMWR conceived the experimental design, carried out data collection, carried out data analyses and drafted the manuscript. ANG participated in data collection. RCR performed phenolic analysis. BDM participated in the design of the study. SNJ participated in the design of the study and in drafting the manuscript. All authors provided feedback on the final manuscript draft and gave approval for publication.

## Data availability

Data will be available at the dryad digital repository upon acceptance (doi to follow).

